# GBMPurity: A Machine Learning Tool for Estimating Glioblastoma Tumour Purity from Bulk RNA-seq Data

**DOI:** 10.1101/2024.07.11.602650

**Authors:** Morgan P. H. Thomas, Shoaib Ajaib, Georgette Tanner, Andrew J. Bulpitt, Lucy F. Stead

**Affiliations:** School of Computing, University of Leeds, UK; Leeds Institute of Medical Research at St James’s, University of Leeds, Leeds, UK

**Keywords:** Glioblastoma, tumour purity, transcriptomics, deconvolution, tumour microenvironment

## Abstract

**Background:** Glioblastoma (GBM) presents a significant clinical challenge due to its aggressive nature and extensive heterogeneity. Tumour purity, the proportion of malignant cells within a tumour, is an important covariate for understanding the disease, having direct clinical relevance or obscuring signal of the malignant portion in molecular analyses of bulk samples. However, current methods for estimating tumour purity are non-specific, unreliable or technically demanding. Therefore, we aimed to build a reliable and accessible purity estimator for GBM.

**Methods:** We developed GBMPurity, a deep learning model specifically designed to estimate the purity of IDH-wildtype primary GBM from bulk RNA-seq data. The model was trained using simulated pseudobulk tumours of known purity from labelled single-cell data acquired from the GBmap resource. The performance of GBMPurity was evaluated and compared to several existing tools using independent datasets.

**Results:** GBMPurity outperformed existing tools, achieving a mean absolute error of 0.15 and a concordance correlation coefficient of 0.88 on validation datasets. We demonstrate the utility of GBMPurity through inference on bulk RNA-seq samples and reveal reduced purity of the Proneural molecular subtype attributed to increased presence of healthy brain cells.

**Conclusions:** GBMPurity provides a reliable and accessible tool for estimating tumour purity from bulk RNA-seq data, enhancing the interpretation of bulk RNA-seq data and offering valuable insights into GBM biology. To facilitate the use of this tool by the wider research community, GBMPurity is available as a web-based tool at: https://gbmdeconvoluter.leeds.ac.uk/.

**Key Points:**

- GBMPurity is a glioblastoma-specific purity estimation tool.
- The model accurately estimates the purity of bulk RNA-seq data, outperforming existing tools.
- The model is available online at: https://gbmdeconvoluter.leeds.ac.uk/.

**Importance of the Study:** Glioblastoma (GBM) is a deadly brain tumour with a dismal prognosis. Research on this disease has lagged compared to other cancers, underscoring the need to streamline investigations. The cellular composition of the GBM tumour microenvironment significantly influences therapy resistance, prognosis, and the molecular state of neoplastic cells. Consequently, tumour purity (the proportion of malignant cells within a tumour) is a critical variable for understanding and contextualizing molecular and clinical analyses. We present GBMPurity (https://gbmdeconvoluter.leeds.ac.uk/), an accessible, GBM-specific tool that accurately predicts sample purity from bulk RNA-seq data. This tool can be used by the wider research community to support the interpretation of bulk omics data and accelerate the identification of more effective therapeutic strategies for treating GBM.

## Introduction

Glioblastoma (GBM) is the most common and aggressive primary brain tumour in adults, with a median survival time between 10 and 14 months and only 30% of patients surviving beyond one year (Mohammed et al., 2022). This dismal prognosis is due to GBM’s rapid growth and diffuse nature combined with a lack of therapeutic innovation, with no significant advancements in treatment strategies for two decades (Stupp et al., 2005).

The complexity of GBM, characterized by substantial inter- and intra-patient heterogeneity in genetic, epigenetic, and cellular landscapes, poses significant challenges to understanding the disease and identifying consistent therapeutic targets (Verhaak et al., 2010; Patel et al., 2014; Neftel et al., 2019; Tanner et al., 2024). Moreover, brain tumour research has been historically underfunded, resulting in slower progress compared to other cancers (Purshouse et al., 2024). This underscores the urgent need to streamline GBM research to better understand its biology and develop more effective treatments.

The GBM tumour microenvironment (TME) is highly heterogeneous and has a well-described effect on malignant progression. It includes a variety of non-malignant cells such as neurons, astrocytes, oligodendrocytes, microglia, infiltrating immune cells, and vasculature. These components influence the state and evolution of GBM cells and contribute to the malignant phenotype, radiotherapy resistance, and overall prognosis (Wang et al., 2017; Martinez-Lage et al., 2019; Pine et al., 2020; Wang et al., 2022; Sharma et al., 2023).

Bulk omics data provides a composite view of all cells within a sample, making tumour purity – a measure of the ratio of malignant to non-malignant cells – a critical factor in data interpretation. Low tumour purity can obscure meaningful signals from the malignant cell fraction, complicating genomic analysis and masking clinical insights (Aran et al., 2015; Haider et al., 2020). Single-cell approaches can overcome this issue, but they are technically challenging and costly, limiting their application to sufficient tissue sample numbers for ascertaining biological and clinical insights in this heterogeneous disease. Consequently, accurately quantifying the contribution of malignant cells to bulk omics data serves as a crucial covariate for deciphering malignant-cell-intrinsic biology.

The purity of a bulk tumour sample can be quantified prior to any molecular analysis by pathology assessment, but these estimates can vary significantly (Smits et al., 2014). Genomic-based methods, which compare somatic copy-number alterations between malignant and non-malignant cell components, offer an alternative. However, low tumour purity combined with high chromosome copy numbers, or high tumour purity with low chromosome copy numbers, can yield similar estimates of purity, resulting in unreliable measurements (Carter et al., 2012; Tanner et al., 2021). This challenge is further complicated by subclonal heterogeneity, for which GBM is notorious (Neftel et al., 2019). Moreover, neither of these methods allows purity inference on publicly available bulk RNA-seq datasets where matched DNA sequencing is not available.

Therefore, RNA-based purity prediction methods are needed. Revkov *et al*. (2023) recently developed a pan-cancer purity estimation tool, PUREE, based on consensus genomic-derived purity labels. However, as previously mentioned, genomic-based estimations may be unreliable. Alternatively, purity estimation can be framed as a cellular deconvolution problem with two cell types, malignant and non-malignant. While there exist multiple RNA-seq compositional deconvolution tools such as CIBERSORTx (Newman et al., 2019), MuSiC (X. Wang et al., 2019), and Scaden (Menden et al., 2020), applying these deconvolution tools can be time-consuming and challenging, particularly for bioinformatics-naïve investigators. Moreover, our previous research (Ajaib et al., 2023) shows that tissue and disease state agnostic deconvolution tools are less accurate than those designed for specific use cases, which is likely to also be true for purity estimation.

To test this, we aimed to build and optimise a tissue-specific purity estimator for GBM, a cancer where purity is particularly pertinent for understanding molecular disease states (Tanner et al., 2024). Our approach leverages deep learning combined with *ad-hoc* simulation of pseudobulked single-cell samples with known purity to augment the amount of training data.

We demonstrate that our approach outperformed general tools: PUREE, CIBERSORTx, MuSiC, and Scaden. Therefore, we developed the model into a ’plug-and-play’ web-based tool called GBMPurity, which is freely available at https://gbmdeconvoluter.leeds.ac.uk/. Users simply upload raw count data of bulk RNA-seq GBM samples and receive accurate estimations of sample purity.

## Materials and Methods

### Data Acquisition and Preprocessing

#### Bulk

Raw, longitudinally matched glioblastoma (GBM) tissue samples were acquired from various sources, with bulk RNA sequencing (RNA-seq) performed according to the protocol described by Tanner *et al*. (2024). We acquired additional raw RNA-seq data from several published studies, following Data Transfer Agreements where required (Kim et al., 2015; Wang et al., 2016; Körber et al., 2019; Kim et al., 2020; Wang et al., 2021; Varn et al., 2022; Hoogstrate et al., 2023). The resulting FASTQ files were processed into a count matrix using the pipeline detailed by Tanner *et al*. (2024). Additionally, we obtained pre-processed RNA-seq data, in the form of transcript counts, from the GLASS consortium via their portal (https://www.synapse.org/glass) (Varn et al., 2022). Information on the availability of these datasets can be found in Table 1.

**Table 1.**
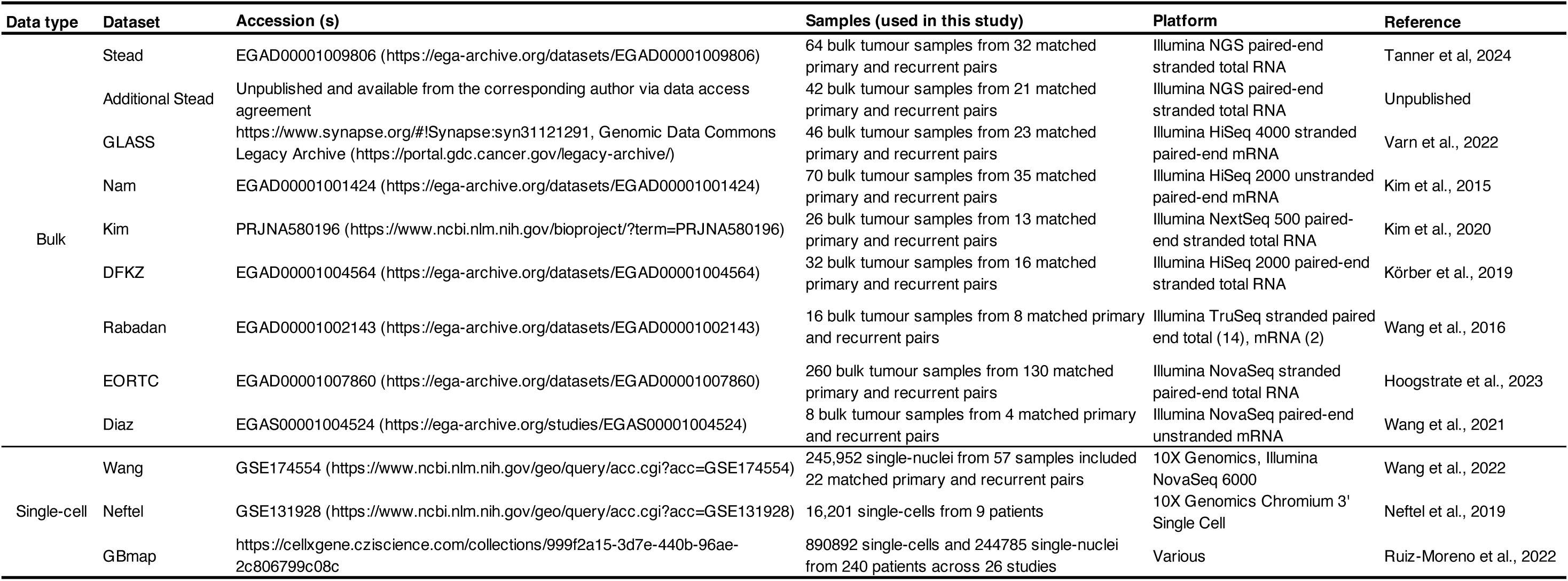
Glioblastoma datasets used in this article.

For the molecular subtyping of bulk tumours, a matrix of transcript per million (TPM) normalized protein-coding genes was uploaded to the GlioVis web application (http://gliovis.bioinfo.cnio.es/) (Bowman et al., 2017). The tumours were then classified according to the consensus of the 3-way SubtypeME tool.

The TPM normalization was performed using the following equation:

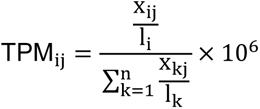

Where *x*_*ij*_ is the raw count of the *i*^th^ gene in the *j*^th^ sample, and *l*_*i*_ is the length of the *i*^th^ gene.

For the deconvolution of bulk samples, the TPM normalized protein-coding genes were uploaded to the GBMDeconvoluteR web application (https://gbmdeconvoluter.leeds.ac.uk/) (Ajaib et al., 2023). The deconvolution was performed using the Ruiz-Moreno marker gene list.

#### Training scRNA-seq data

The training single-cell RNA-seq (scRNA-seq) dataset was obtained from the extended <u>GBmap resource</u>, an integrated and annotated single-cell atlas of IDH-wildtype primary GBM encompassing over 1.1 million cells from 240 patients across 26 studies (Ruiz-Moreno et al., 2022). These data were pre-filtered by the original authors for cells that expressed over 500 genes, 1000 RNA counts and less than 30% mitochondrial reads.

#### Validation scRNA-seq data

Two validation IDH-wildtype primary GBM datasets were utilized. The first, single-nucleus RNA-seq data (n = 57), was downloaded from GSE174554 (Wang et al., 2022) and processed using Seurat’s (v5.1.0) SCTransform method. Malignant cells in this dataset were pre-annotated by the original authors.

The second scRNA-seq dataset was retrieved from GSE131928 (Neftel et al., 2019). This dataset comprises a mixture of Smart-seq2 and 10X single-cell profiling data. Given that the Smart-seq2 data were enriched for malignant cells, only the 10X data were included in this study to ensure a range of purities for model evaluation (n = 9). As this dataset is part of GBmap, samples used for validation were excluded from the GBmap dataset to avoid data leakage. This data was filtered using Seurat (v5.1.0) for cells with more than 800 RNA counts, over 200 detected genes, and less than 5% mitochondrial gene content. Doublets were identified and removed using the DoubletFinder package (v2.0.4) (McGinnis *et al*., 2019).

Malignant cells were identified through copy number alteration (CNA) analysis using the CONICSmat package (v0.0.0.1) (Müller et al., 2018) (Supplementary Fig. 1a). Cells were annotated as malignant if they had a posterior probability greater than 0.95 of harbouring one of the known GBM chromosomal aberrations: Chromosome 7 arm p and q amplification or Chromosome 10 arm q deletion (Supplementary Fig. 1d). The single-cell data were then integrated using Seurat’s (v5.1.0) IntegrateLayers with the CCAIntegration method after applying the NormalizeData, FindVariableFeatures, and RunPCA methods with default parameters (Supplementary Fig. 1b, c).

### GBMPurity Model Development

#### Feature selection

To develop the GBMPurity model aimed at inferring the purity of bulk RNA-seq GBM samples using scRNA-seq data, our feature selection process focused on identifying genes that are consistently represented across both RNA-seq modalities. We compared bulk RNA-seq data to the GBmap single-cell data after pseudobulking, which involved summing RNA-seq counts of cells from the same sample. We first excluded genes that were either absent or expressed at low levels (counts per million (CPM) < 1 in 50% of samples) in either modality.

The CPM normalization was performed using the following equation:

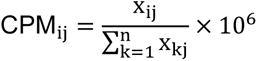

Where *x*_*ij*_ is the raw count of the *i*^th^ gene in the *j*^th^ sample.

Following CPM normalization, we employed the Kolmogorov-Smirnov (KS) statistic to quantify the distance between the empirical distribution functions of each gene in the two modalities, providing a measure of similarity between the distributions. Genes with a KS statistic above an arbitrary threshold of 0.4 were excluded, resulting in a final set of 5829 genes for model training. A visual representation of this filtering process is provided in Supplementary Fig. 2.

#### Simulation of pseudobulk samples

Deep learning models typically contain a large number of learnable parameters, enabling them to capture highly complex functions. However, this also makes them prone to overfitting in the absence of extensive training data. To mitigate this risk, we simulated pseudobulk tumours during the training process, which allowed us to generate sufficient training data. We preserve interpatient heterogeneity by sampling cells only from within the same GBM sample. Samples containing fewer than five malignant or non-malignant cells were excluded to ensure an adequate number of cells for sampling across the purity spectrum.

To simulate pseudobulk tumours for a given sample *i*, two random numbers are generated: the target purity *p* (0 to 1), and the number of cells *N* (200 to 4000). Based on *p* and *N*, the number of malignant (*N*_*m*_) and non-malignant (*N*_*n*_) cells are calculated. Cells (*x*) are randomly sampled with replacement from the selected sample. The RNA-seq counts are summed across the sampled malignant and non-malignant cells to simulate a pseudobulk tumour:

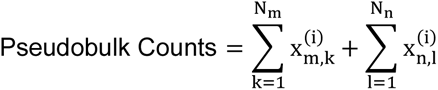

#### Input data processing

Simulated pseudobulk samples undergo several transformation steps to prepare the data for model training. The steps are as follows:

1. The raw counts of the simulated pseudobulk samples are first TPM transformed.
2. The TPM values are then divided by 100 to rescale the data to a more suitable range for model training.
3. A log_2_ transformation is applied to the scaled TPM values after adding 1 to each value to avoid infinite values:

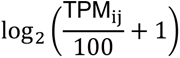

#### Model construction

GBMPurity is a regression machine learning model developed using PyTorch (v2.2.0), designed to predict the purity of bulk GBM samples from an input of 5829 selected genes. The model outputs a single numeric value representing the estimated purity. It was trained using the Adam optimizer, processing data in batches of 64 randomly simulated pseudobulks until convergence of the L1 loss function. Convergence was defined as the point where the average training loss failed to decrease over a sliding window of 25 batches.

L1 loss is defined as:

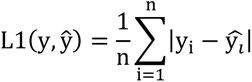

Where *y*_*i*_ is the actual purity of *i*^th^ sample and *ŷ*_*i*_ is the predicted purity.

The model’s performance relies on several hyperparameters which were optimized through cross-validation. Each dataset in the GBmap resource was treated as an individual fold. For effective evaluation across a representative spectrum of purity, we focused on 11 of the 26 folds that contained at least five samples with purity values between 0.1 and 0.9. These 11 folds were used as holdout datasets, resulting in 11 cross-validation iterations.

The following hyperparameters were tuned sequentially in independent experiments (as shown in Supplementary Figure 3): the number of hidden layers, the size of the hidden layer dimensions, the dropout rate, weight decay, learning rate, and patience (defined as the number of batches to wait for a decrease in loss before terminating training). These hyperparameters were fine-tuned to achieve optimal model performance, ensuring robustness and generalizability in predicting the purity of bulk GBM samples.

#### GBMPurity model

The resulting GBMPurity model is a multi-layer perceptron with two hidden linear layers, comprising 32 and 16 neurons respectively, each employing a rectified linear unit (ReLU) activation function. The model was trained with a learning rate of 3e^-5^, a weight decay of 1e^-5^, and an input layer dropout probability of 0.4. We saved the model with the lowest average loss over a 25-batch sliding window and terminated training when this sliding average did not decrease for 200 batches.

During inference, the 5829 selected genes are input into the model. The model outputs a continuous prediction of purity, which can theoretically range from negative infinity to infinity due to the absence of an activation function on the output layer. To ensure meaningful predictions, we manually clip these outputs to a range between 0 and 1.

#### Model evaluation

For the evaluation of GBMPurity, along with other benchmarked models described below, we input pseudobulks of true samples from Wang *et al*. (2022) and Neftel *et al*. (2019) and measure the error of the predictions versus the labelled true purity. We describe the performance across four metrics: mean absolute error (MAE), root mean squared error (RMSE), Pearson correlation, and Correlation Concordance Correlation (CCC):

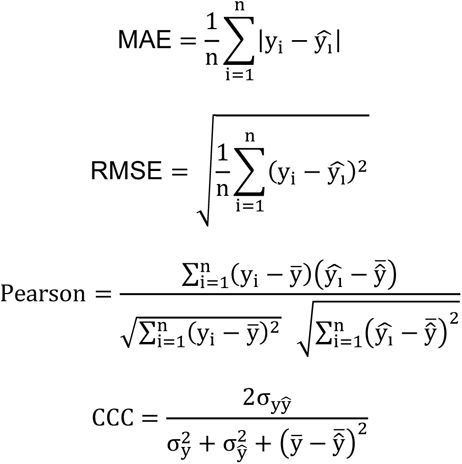

Where *y*_*i*_ is the actual value and *ŷ*_*i*_ is the predicted value of the *i*^th^ sample, and *n* is the number of observations.

### Model Interpretation

To understand and validate the predictions of GBMPurity, we employed techniques for feature attribution and model interpretation.

#### SHapley Additive exPlanations (SHAP)

We utilized SHAP to determine each feature’s impact on the predicted purity in the pseudobulked training data (Lundberg and Lee, 2017). This was implemented using the DeepExplainer class from the Python shap package (v0.45.1). SHAP values provide a measure of each gene’s contribution to the model’s output, allowing for a detailed understanding of feature importance.

#### Gene Set Enrichment Analysis (GSEA)

Following SHAP, we generated a pre-ranked list by averaging the SHAP contributions for each feature across the training data. Gene Set Enrichment Analysis (GSEA) was then performed using the brain-related gene set database curated by Hagenauer *et al*. (2024) with the fgsea package (v1.26.0) in R (Korotkevich et al., 2021). This analysis was conducted without ranking metric weighting.

#### Interpretation of hidden nodes

To further interpret the internal workings of the model, we used the LayerConductance class from the Python package Captum (v0.7.0) to quantify the average contribution of each hidden neuron to the model output (Dhamdhere et al., 2018).

### Benchmarking

We compared the performance of GBMPurity against four established tools that enable purity prediction. Where single-cell references were necessary for training the model, we used the GBmap dataset with the same 5829 genes and cells labelled as malignant or non-malignant, mirroring the training process of GBMPurity. The predicted contribution of the malignant component was taken as the model’s prediction of tumour purity. Each model was evaluated using MAE, RMSE, Pearson correlation, and CCC.

#### PUREE

PUREE (Revkov et al., 2023) is a pan-cancer purity estimation tool that employs a weakly supervised machine learning model trained on RNA-seq data across multiple cancer types labelled with consensus purity estimates derived from four different algorithms. This pre-trained model did not require single-cell reference data. For purity estimation, pseudobulks of the three single-cell datasets used in this study were TPM transformed and uploaded to the PUREE web interface (https://puree.genome.sg/).

#### CIBERSORTx

CIBERSORTx (Newman et al., 2019) is a cell deconvolution algorithm that uses single-cell reference data to generate gene expression profiles of various cell types. It employs support vector regression to estimate the proportions of different cell types in an RNA-seq mixture. Due to the size of our single-cell reference, CIBERSORTx was run via Docker and used the default parameters.

#### MuSiC

MuSiC (X. Wang et al., 2019) deconvolves bulk RNA-seq data using similar means to CIBERSORTx, but instead utilizes sample information to weight genes with consistent cross-subject and cross-cell type consistency, employing a non-negative least squares algorithm. The deconvolution was performed in R using the music_prop function from the MuSiC package (v1.0.0) with default parameters.

#### Scaden

Scaden (Menden et al., 2020) is an ensemble deep-learning model that deconvolves bulk RNA-seq samples using labelled pseudobulks. This approach is similar to the methods employed in GBMPurity; however, Scaden uses pre-simulated pseudobulks of fixed cell numbers and fixed training steps, whereas GBMPurity generates pseudobulk samples of varying cell numbers during training and terminates training automatically when the loss stops decreasing. We trained Scaden using default parameters on 500 simulated tumours per training sample, each containing 500 cells.

### Statistical Analysis

All statistical analyses were conducted using Python, specifically leveraging the pingouin package (v0.5.4) for statistical computations. Descriptive statistics, inferential tests, and correlation analyses were performed to validate the findings. P-values less than 0.05 were considered statistically significant.

### Software and Hardware use

All computational analyses were performed using ARC4, part of the High-Performance Computing facilities at the University of Leeds, UK. This system runs the CentOS 7 distribution of Linux and contains Intel Xeon Gold 6138 CPUs with up to 768GB of memory. All analyses were conducted in R version 4.3.1 or Python version 3.10.13.

### Data and Code availability

The datasets generated and analysed during the current study are available as described in the original publications. All code used for data processing, model development, and analysis is available at https://github.com/scmpht/GBMPurity. The pre-trained GBMPurity model, along with instructions for use, is available at https://gbmdeconvoluter.leeds.ac.uk/.

### Ethics Statement

All data used in this study was derived from patients who provided samples with informed, written consent. These samples were approved for use in this study by the UK National Health Service’s Research Ethics Service Committee South Central – Oxford A (Research Ethics Code: 13/SC/0509).

## Results

### Data Collection and Preprocessing

To develop GBMPurity, GBmap, a comprehensive glioblastoma (GBM) single-cell RNA-seq atlas curated by Ruiz-Moreno *et al*. (2022), was utilized. This dataset comprises integrated single-cell RNA-seq data from 240 GBM patients across 26 studies. Additional validation datasets were obtained from Wang *et al*. (2022), consisting of 57 pre-labelled single-cell GBM samples, and Neftel *et al*. (2019), including 9 unlabelled samples that were manually labelled (Supplementary Fig. 1). The Neftel samples were also included in GBmap and thus excluded from the training set, resulting in 231 training and 66 validation samples (Fig. 1a).

**Figure 1.**
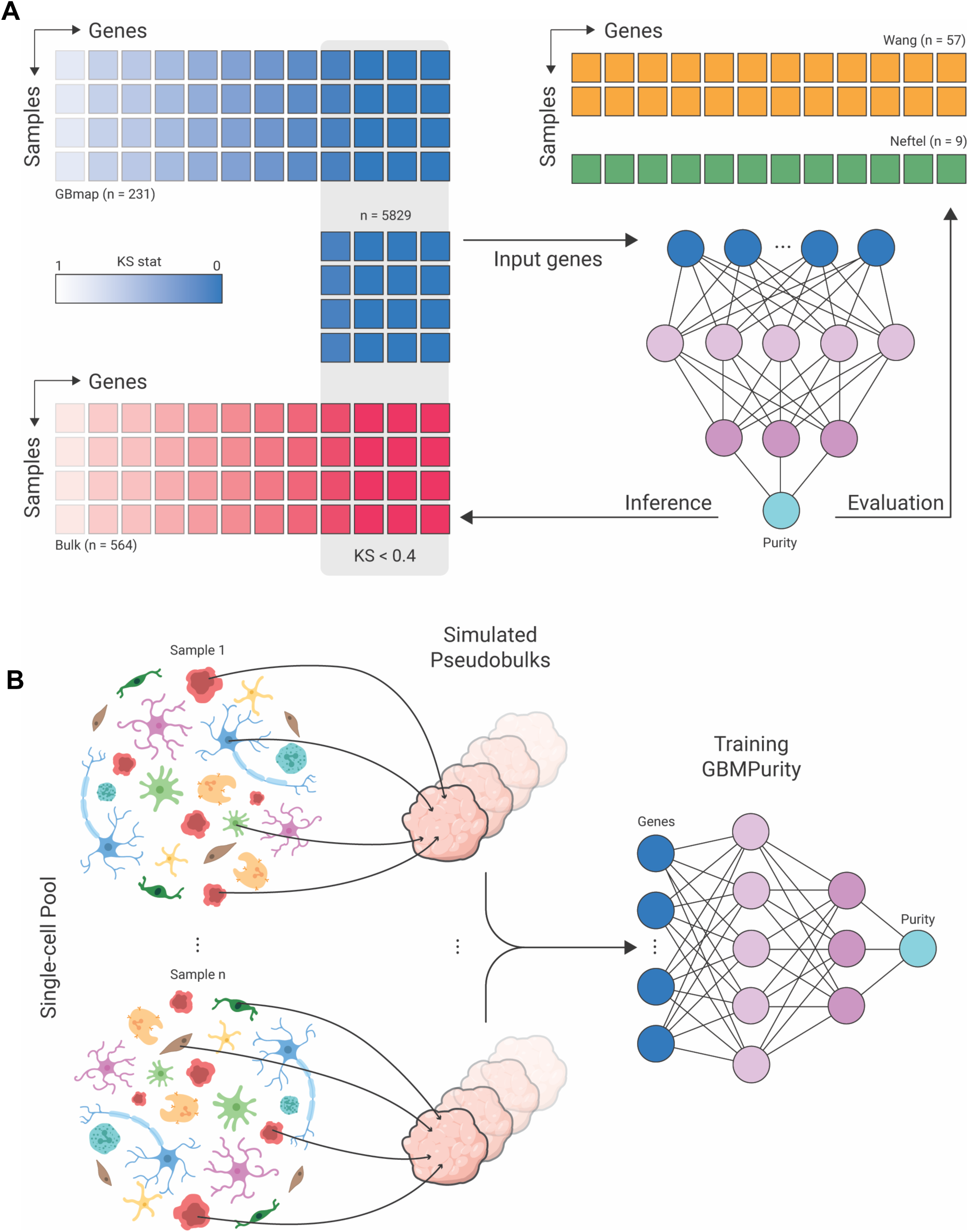
Study design and training methodology for GBMPurity. (a) Single-cell RNA-seq data from the GBmap dataset (Ruiz-Moreno et al., 2022) was compared to in-house bulk GBM RNA-seq data (n = 564) to identify genes with a Kolmogorov-Smirnov statistic (KS stat) of < 0.4 between the two modalities. These genes were used to train the GBMPurity model. The trained model was evaluated on 57 pre-labelled single-cell pseudobulks from Wang *et al*. (2022) and 9 from Neftel *et al*. (2019), which were labelled using CONICSmat (Müller *et al*., 2018). The model was subsequently applied to bulk RNA-seq data for purity inference. (b) Random sampling of cells from within-patient samples was used to simulate pseudobulks of known purity. These simulated samples were used to train GBMPurity until convergence of the L1 loss function.

To ensure the generalizability of our model to bulk RNA-seq samples, we selected genes equally represented across the pseudobulked single-cell GBmap samples and the bulk GBM samples. Following counts per million (CPM) normalization, genes with CPM < 1 in over 50% of samples in either modality were excluded. Using a Kolmogorov-Smirnov statistic threshold of < 0.4, we selected 5,829 genes with similar distributions across both modalities for model training (Fig. 1a, Supplementary Fig. 2).

### Model Development

We simulated pseudobulk samples with known purity by random sampling of malignant and non-malignant cells, ensuring that pseudobulks were kept within-sample to maintain robustness against intratumoural heterogeneity (Fig. 1b). Samples with fewer than five malignant or non-malignant cells were excluded, resulting in 197 samples used for simulation. Pseudobulks were simulated *ad hoc* during model training until the mean absolute error (MAE) loss function converged. Hyperparameters were optimized using cross-validation (Supplementary Fig. 3), and the final model, which we named GBMPurity, was trained using all 197 samples.

### Model Evaluation

GBMPurity demonstrated robust performance across multiple validation datasets, depicted in Figure 2a and detailed in Table 2. Specifically, the model achieved a Mean Absolute Error (MAE) of 0.15 on both the Wang and Neftel data, with a CCC of 0.88 and 0.77, respectively, demonstrating the high accuracy of our model.

**Figure 2.**
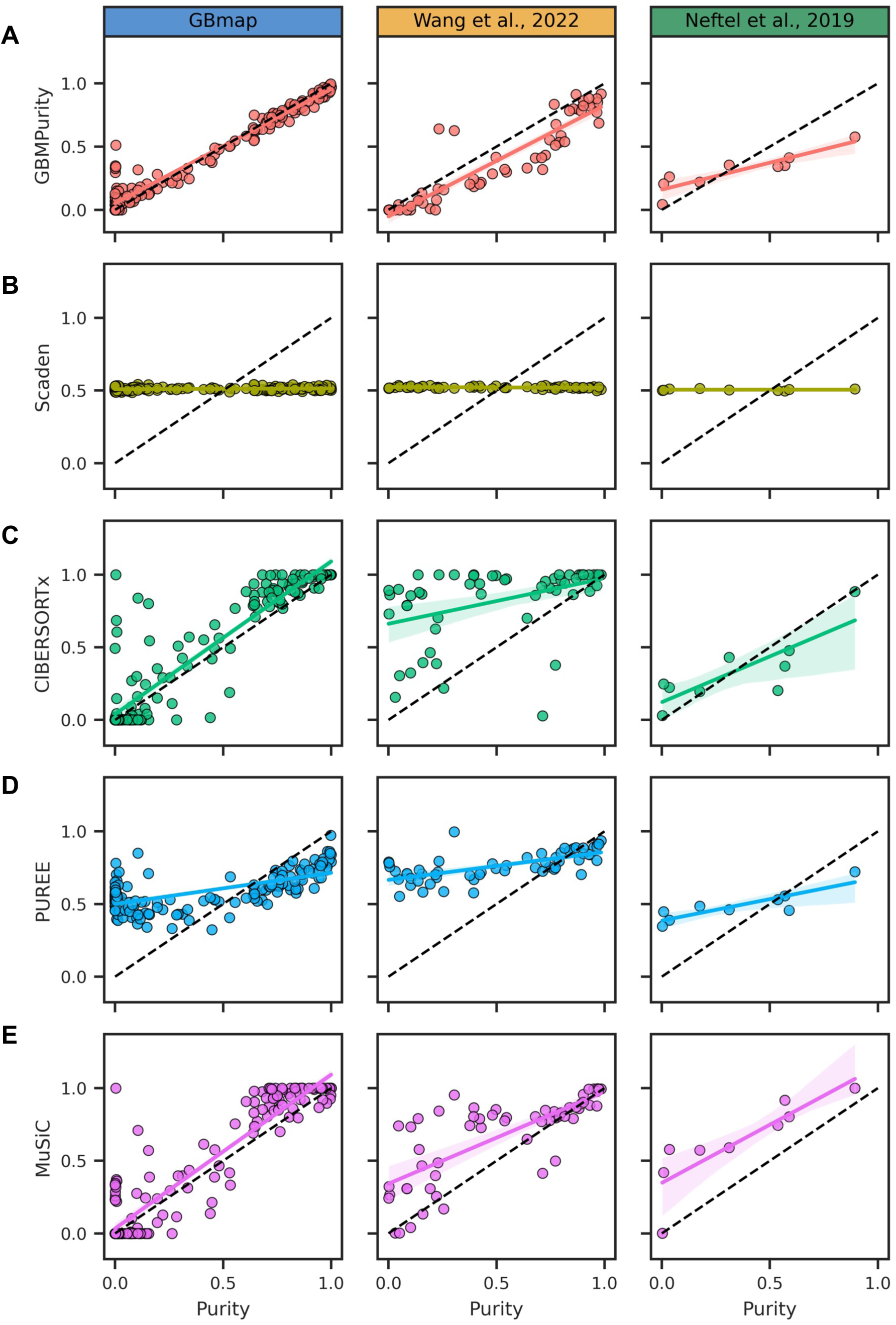
Benchmarking of GBMPurity against alternative purity estimation methods. Each panel presents the relationship between true and predicted purity across three datasets; GBMap, Wang *et al*., 2022, and Neftel *et al*. 2019. The models compared are (a) GBMPurity, (b) Scaden, (c) CIBERSORTx*, (d) PUREE, and (e) MuSiC. GBmap was used as training data for all models requiring reference data. Performance metrics are detailed in Table 1.

**Table 2.** Purity estimation benchmarking results.

| Dataset | Model | MAE | RMSE | Pearson's | CCC |
| --- | --- | --- | --- | --- | --- |
| GBmap | GBMPurity | **0.046** | **0.083** | **0.978** | **0.980** |
|  | Scaden | 0.377 | 0.401 | 0.188 | 0.010 |
|  | CIBERSORTx | 0.096 | 0.172 | 0.929 | 0.917 |
|  | PUREE | 0.315 | 0.372 | 0.667 | 0.313 |
|  | MuSiC | 0.105 | 0.163 | 0.937 | 0.926 |
| Wang et al., 2022 | GBMPurity | **0.145** | **0.177** | **0.917** | **0.879** |
|  | Scaden | 0.310 | 0.339 | -0.287 | -0.013 |
|  | CIBERSORTx | 0.347 | 0.445 | 0.422 | 0.269 |
|  | PUREE | 0.283 | 0.375 | 0.667 | 0.241 |
|  | MuSiC | 0.189 | 0.271 | 0.751 | 0.674 |
| Neftel et al., 2019 | GBMPurity | 0.162 | 0.186 | **0.902** | 0.773 |
|  | Scaden | 0.286 | 0.339 | -0.003 | 0.000 |
|  | CIBERSORTx | **0.139** | **0.174** | 0.815 | **0.887** |
|  | PUREE | 0.214 | 0.259 | 0.864 | 0.496 |
|  | MuSiC | 0.279 | 0.320 | 0.854 | 0.656 |

MAE, mean absolute error; RMSE, root mean squared error; CCC, correlation concordance coefficient.

We then investigated the robustness of GBMPurity. A tendency to underestimate purity was observed in the validation data (Supplementary Fig. 4a). This observation likely reflects the increased stringency of malignant cell assignment during the labelling process between the training and validation datasets (Supplementary Fig. 4b), rather than a bias in the model predictions. Notably, the model performed as expected when using the excluded Neftel samples from the GBmap dataset (Supplementary Fig. 4c).

The robustness of GBMPurity to missing genes was also evaluated, anticipating that not all datasets would contain a complete gene set. The model maintained high performance with 10-20% missing genes highlighting its resilience to varying data completeness (Supplementary Fig. 4d, e). Lastly, we evaluated the stability of the model by training GBMPurity with different weight initialisations and observed consistency in model performance across the three datasets (Supplementary Fig. 4f).

### Benchmarking

We evaluated GBMPurity’s performance against established purity estimation tools. PUREE (Revkov et al., 2023), which demonstrated good performance against an extended set of methods, served as our reference-free benchmark. We also included CIBERSORTx (Newman et al., 2019) since it is the most established of the single-cell reference deconvolution tools, and MuSiC (X. Wang et al., 2019) as it is another popular reference-based method that wasn’t included in PUREE’s benchmarking. Lastly, Scaden (Menden et al., 2020) another machine-learning model trained on simulated pseudobulks, was included because of the similarity of methods used to train GBMPurity.

MAE, RMSE, Pearson correlation and CCC performance metrics were assessed across three datasets (Fig. 2, Table 2). GBMPurity consistently outperformed all other tools across nearly all evaluation metrics in the validation datasets, demonstrating the model’s accuracy and robustness in estimating tumour purity from RNA-seq data.

### Model Interpretation

Despite the inherent complexity of deep learning models, we sought to interpret GBMPurity to validate its predictions and derive biological insights. We applied SHAP (Lundberg and Lee, 2017) – which quantifies the importance of each input feature for a prediction – to the pseudobulks of the training GBmap dataset (Fig. 3a). The use of dropout and weight decay to prevent overfitting resulted in a distribution of small impacts across many genes. MT-RNR2 like 12 (*MTRNR2L12)* and MT-RNR2 like 8 (*MTRNR2L8)* emerged as the most influential features, both contributing to higher purity estimates. As pseudogenes, the literature on these species is sparse, but their function is estimated to be involved in the negative regulation of apoptosis, a hallmark of cancer (Bult and Sternberg, 2023). Cysteine Rich Protein 1 (*CRIP1)*, which is predominantly expressed in blood and immune cells (Karlsson et al., 2021), was the most influential gene associated with lower purity estimates.

**Figure 3.**
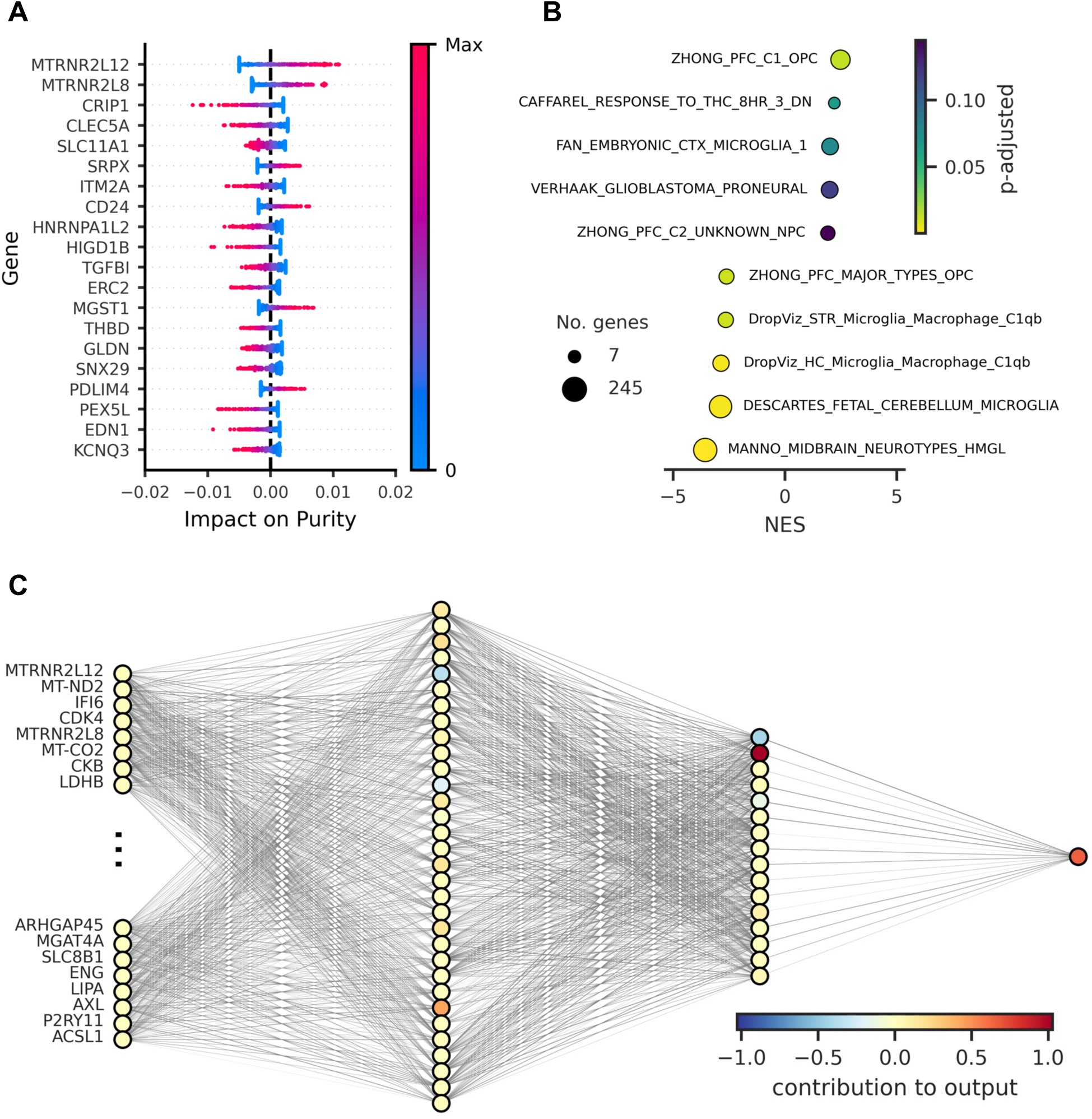
Interpretation and architecture analysis of GBMpurity. (a) SHapley Additive exPlanations (SHAP) summary plot of the top 20 most influential features on model predictions in the pseudobulked GBmap data. (b) Gene set enrichment analysis of genes ranked by their average SHAP value across the pseudobulked GBmap data using a brain-specific gene set database (Hagenauer *et al*., 2024). (c) Visualisation of GBMPurity model architecture with neurons coloured by average contribution to purity estimation determined with conductance analysis (Dhamdhere *et al*., 2018).

We then ranked genes based on the average magnitude of their SHAP contributions over the pseudobulked training data, emphasizing broad average impact rather than rare, high-magnitude impacts. This ranking was used for GSEA with a curated database of brain-related gene sets (Hagenauer et al., 2024). GSEA identified that genes associated with neurodevelopmental precursors positively influenced purity estimates, while microglia and neuronal gene sets negatively impacted purity estimates (Fig. 3b). These results instil confidence in our model as single-cell studies have shown that neoplastic GBM cells hijack neurodevelopmental processes (Couturier et al., 2020), and microglia, as resident brain macrophages, along with neurons, the primary brain cell type, are key components of the non-neoplastic GBM microenvironment.

Conductance analysis was employed to quantify the importance of each node within the hidden layers (Dhamdhere et al., 2018). Visualization of GBMPurity and the importance of each node is shown in Figure 3c. This method can also be used to quantify the contribution of each input feature (i.e. expression of associated genes) to specific nodes. GSEA of these rankings suggested these nodes are polysemantic, not representing distinct identifiable biological modules (data not shown).

### Bulk Inference

Having established GBMPurity’s efficacy in estimating the purity of pseudobulked single-cell RNA-seq data, we used it to infer the purity of bulk RNA-seq samples. The raw count matrix from the bulk RNA-seq data was processed through the GBMPurity model, which involves automatic filtering and transformation of the input data before being fed through the neural network. The model subsequently outputs a list of estimated purities, each corresponding to a specific sample in the dataset.

To validate the inferences of GBMPurity against established biological knowledge, we inspected purity across various biological subgroupings of GBM. Firstly, it is well-documented that tumour purity tends to decrease upon recurrence (Wang et al., 2022; Hoogstrate et al., 2023) so we compared the purity estimates of primary and recurrent GBM tumours. In validation of our method, our analysis revealed a significant reduction in tumour purity following standard treatment protocols (Fig. 4a).

**Figure 4.**
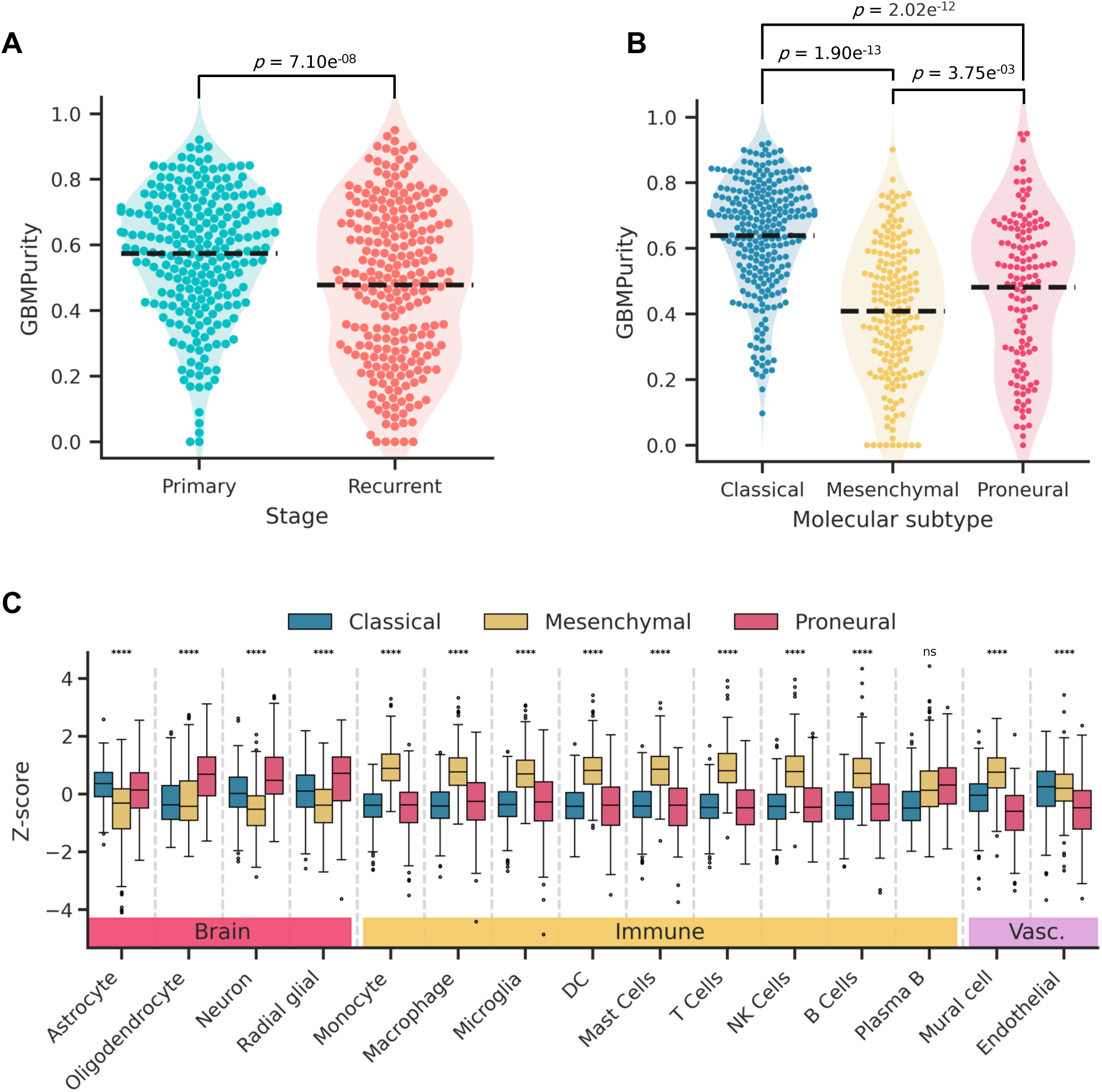
Proneural tumours exhibit a reduction in purity driven by increased normal brain cells. (a, b) Beeswarm plots of GBMPurity estimates of bulk IDH^wt^ primary GBM tumours stratified by (a) stage (n = 482, filtered for patients with local recurrence who underwent the Stupp treatment protocol), and (b) GBM molecular subtype (n = 564). Dashed lines represent the sample means. (c) Boxplot of GBMDeconvoluteR scores (Ajaib *et al*., 2023) for the bulk IDH^wt^ primary GBM tumours stratified by molecular subtype. Significance in (a) was derived using a paired t-test; in (b), an ANOVA followed by pairwise Tukey tests was used. In (c), a mixed ANOVA on the entire dataset was followed by pairwise Tukey tests to assess differences between molecular subtypes within each cell score and subsequent Bonferroni correction. The reported significance level is between Mesenchymal and Proneural cell scores, where **** = p < 0.0001. Full pairwise comparisons can be found in Supplementary Table 1. DC, dendritic cells.

Additionally, GBM tumours can be stratified into Classical, Mesenchymal, and Proneural molecular subtypes, each associated with distinct biological characteristics (Verhaak et al., 2010; Wang et al., 2017). Our model corroborated that Mesenchymal tumours exhibit notably lower purity levels, which is consistent with previous findings (Fig. 4b; Wang et al., 2017; Hoogstrate et al., 2023). However, we also observed a similar reduction in purity in the Proneural subtype, which contrasts with the findings of the aforementioned studies. To investigate this further, we applied GBMDeconvoluteR, a GBM-specific deconvolution tool that provides cell-type-specific scores from bulk RNA-seq data. This analysis revealed that Proneural tumours have a significant increase in normal brain cell types, whereas Mesenchymal tumours have a significant increase in immune cell types (Fig. 4c, Supplementary Table 1).

## Discussion

In this study, we present GBMPurity, a novel deep-learning tool tailored to estimate tumour purity from bulk RNA-seq data, specific to glioblastoma (GBM). We trained a multi-layer perceptron using the extensive single-cell RNA-seq atlas, GBmap (Ruiz-Moreno et al., 2022). This training data was enhanced by simulating pseudobulk samples, enabling the application of a more sophisticated model.

GBMPurity demonstrated robust performance in accurately predicting tumour purity across multiple validation datasets. The model achieved high concordance correlation coefficients (CCC) of 0.88 and 0.77 on the Wang and Neftel datasets, respectively, surpassing established deconvolution tools CIBERSORTx (Newman et al., 2019), MuSiC (X. Wang et al., 2019), PUREE (Revkov et al., 2023), and Scaden (Menden et al., 2020).

GBMPurity offers two main advantages. First, it is a ’plug-and-play’ web-based tool that simplifies the process for GBM researchers, making sophisticated purity estimation accessible even to those with limited bioinformatics expertise. This tool can be applied to pre-existing bulk RNA-seq datasets without requiring extensive computational resources or advanced technical knowledge. Second, the tailoring of this model to GBM has resulted in improved performance compared to general deconvolution and purity estimation methods.

Interpretation of this model also allows us to derive biological inferences. By employing SHAP for model interpretation, we identified key genes influencing tumour purity, providing insights that can guide further biological investigations. Notably, *MTRNR2L12* and *MTRNR2L8* emerged as the most influential genes in purity estimations. These are isoforms of the *MT-RNR2* gene, which encodes Humanin—a small peptide with neuroprotective and anti-apoptotic activity in neuroblasts of Alzheimer’s diseased brains—and has recently been attributed with oncogenic effects in glioblastoma cells (Ying et al., 2004; Bodzioch et al., 2009; Peña Agudelo et al., 2023).

GBMPurity’s inference on bulk RNA-seq samples provided biologically sound insights into the purity of primary versus recurrent tumours, corroborating the known clinical pattern of reduced tumour purity upon recurrence (Wang et al., 2022; Hoogstrate et al., 2023). Additionally, when stratifying GBM samples by molecular subtypes defined by Verhaak, *et al*. (2010) we observed a significantly lower purity in Mesenchymal and Proneural subtypes relative to Classical. While the association of the Mesenchymal subtype with increased immune infiltrate is well documented (Kaffes et al., 2019; Martinez-Lage et al., 2019), the lower purity in Proneural subtypes akin to that of the Mesenchymal subtypes was unexpected as it disagrees with the findings of Hoogstrate *et al*. (2023) who derive purity through CNA-based methods. Deconvolution of these tumours suggests the reduced purity in Proneural tumours can be attributed to the increased presence of normal resident brain cells, as opposed to the increased immune infiltrate of Mesenchymal tumours. This may indicate limitations in the ability of CNA-based purity estimation methods to differentiate malignant cells of Proneural tumours from healthy brain cells. Indeed, we show here a relatively large discrepancy between CNA- and RNA-seq-derived purity estimates (Supplementary Fig. 5a), and no tendency for GBMPurity to underestimate the purity of Proneural tumours (Supplementary Fig. 5b).

The enriched association of, and interaction between, neoplastic GBM cells and normal brain cells within Proneural tumours is now well documented (Venkatesh et al., 2019; L. Wang et al., 2019; Tanner et al., 2024) alongside the repeated observation of Mesenchymal cells being associated with brain resident and infiltrating immune cells (Neftel et al., 2019; Varn et al., 2022). Our results corroborate these findings, further adding confidence to the accuracy of GBMPurity. However, the fact that many studies did not note the reduced purity in Proneural tumours highlights that previous estimates of purity have relied too heavily on proportions of immune cells within the TME, not accounting for the normal brain cell component. This is likely because the latter is harder to distinguish owing to the large overlap in overall expression, and that of marker genes, between normal brain and neoplastic GBM cells. These findings underscore the potential of GBMPurity to enhance the interpretation of bulk RNA-seq data and provide more accurate biological insights into GBM.

Applications of GBMPurity hold promise for advancing glioblastoma research by streamlining the analysis of bulk GBM omics data and offering a more pragmatic interpretation in the context of TME composition. This utility may also extend beyond the lab and into the clinic. For example, immunotherapies are a promising approach in GBM treatment, but as of yet, have produced variable responses (Lim et al., 2018; Agosti et al., 2023). Applying GBMPurity to differentiate between responders and non-responders to immunotherapy could enhance prognostic assessments, given the associations between TME composition and therapeutic response in other solid tumours (Gong et al., 2020; Petitprez et al., 2020).

Future iterations of this model could facilitate automatic correction of the input matrix for estimated purity, enabling more targeted and consistent analyses of the malignant components in bulk omics data. Additionally, while our current approach selected genes from single-cell RNA-seq data that are representative of our bulk RNA-seq dataset, incorporating batch correction methods, such as those implemented by CIBERSORTx (Newman et al., 2019), could further enhance inter-modal applicability. Moreover, varying the ratios of different cell types in the non-malignant component of simulated pseudobulks could improve the diversity of the training data. Finally, our methodology is not restricted to GBM and has the potential to be extended to other cancer types, paving the way for a comprehensive pan-cancer purity estimation tool.

## Acknowledgements

The authors would like to thank the University of Leeds HPC services and CDT in AI for Medical Diagnosis and Care for their support. We would also like to extend our gratitude to the patients and their families who agreed to provide tissue samples that were used in this study and the wider research supporting it.

