## Supplementary Figures for "GBMPurity: A Machine Learning Tool for Estimating Glioblastoma Tumour Purity from Bulk RNA-seq Data"

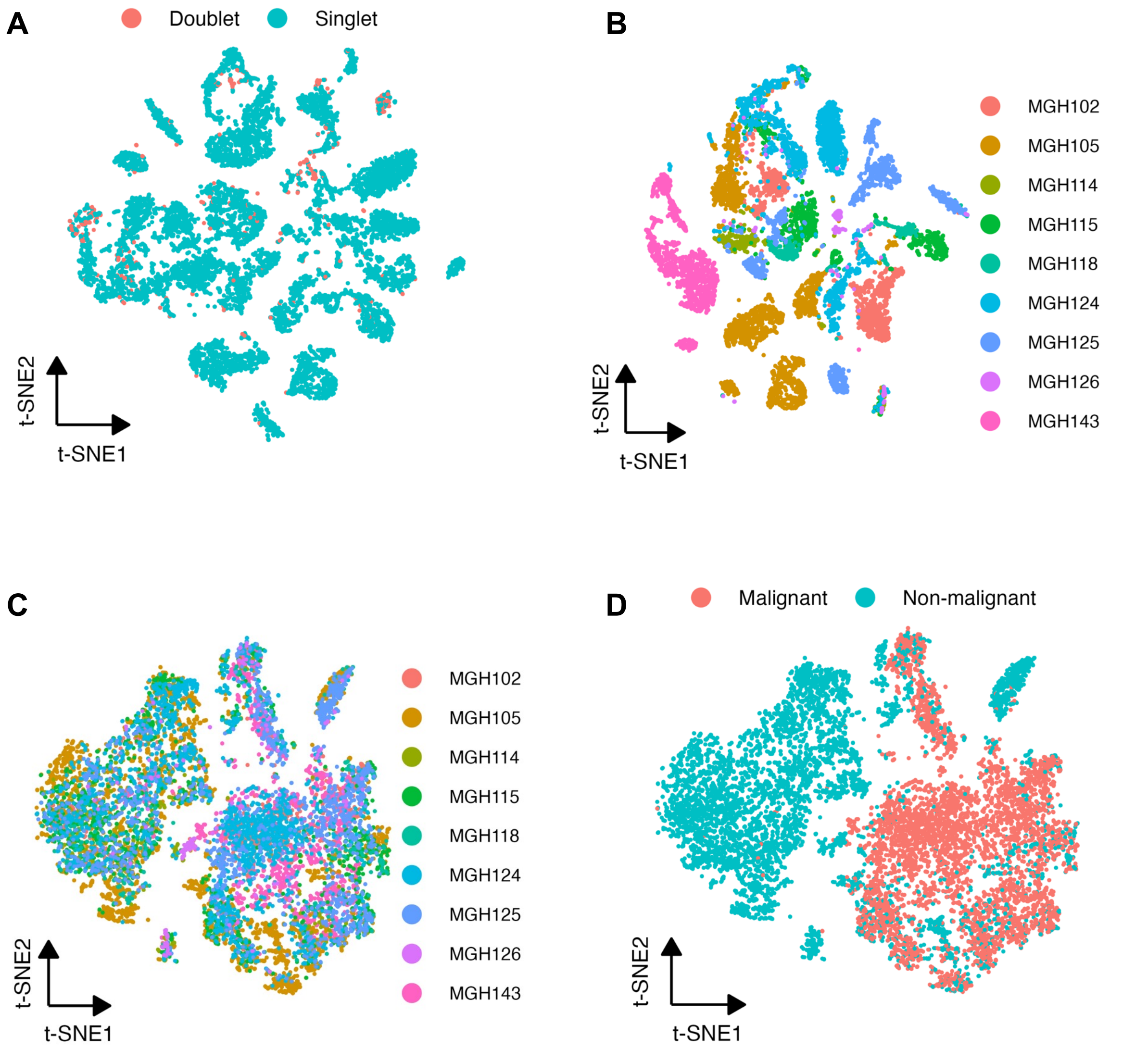


**Supplementary Figure 1. Annotation of malignant cells in the Neftel et al. (2019) dataset.** t-SNE representations of (a) doublets identified and removed with DoubletFinder (McGinnis et al., 2019). (b) before and (c) after integration of samples. (d) Labelling of malignant cells based on copy number alteration (CNA) analysis using CONICSmat (Müller et al., 2018). Cells were annotated as malignant if they had a posterior probability greater than 0.95 of harbouring known GBM chromosomal aberrations.

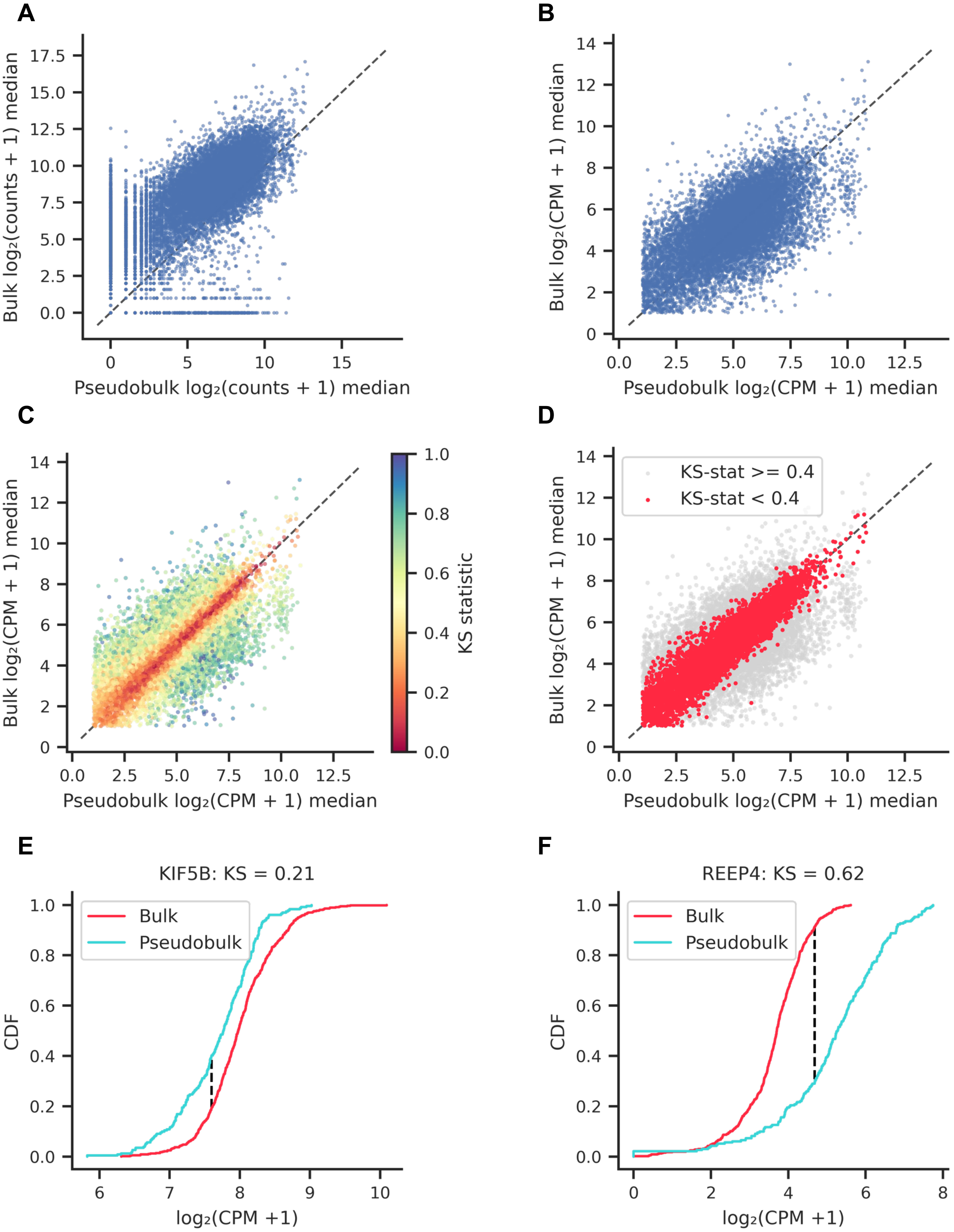


**Supplementary Figure 2. Gene selection for consistent representation across pseudobulk single-cell and bulk modalities.** (a-d) Medians of log2-transformed gene expression values in bulk versus pseudobulk single-cell data during sequential filtration steps. (a) Untransformed data. (b) Counts per million (CPM) transformed and low expression filtered data. (c) Genes coloured by Kolmogorov-Smirnov (KS) statistic. (d) Final set of 5,829 genes with a KS statistic < 0.4 used for training the GBMPurity model. (e, f) Cumulative density function examples illustrating genes with (e) smaller and (f) larger KS statistics.

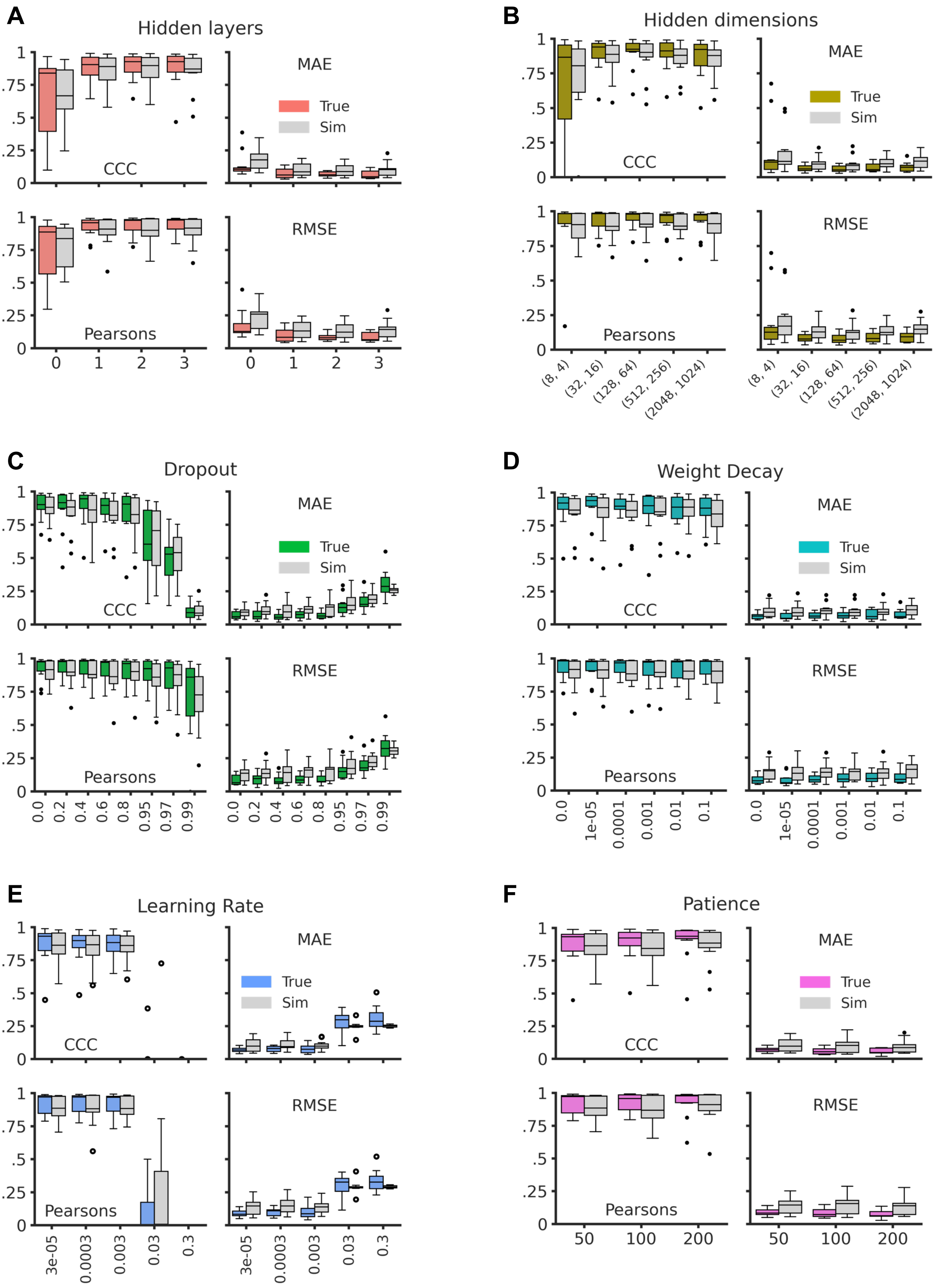


**Supplementary Figure 3. Hyperparameter optimization for the GBMpurity model.** (a-f) Boxplots of model evaluation metrics, including concordance correlation coefficient (CCC), mean absolute error (MAE), Pearson correlation, and root mean squared error (RMSE), for GBMPurity trained with cross-validation across different hyperparameter values. Hyperparameters were tuned sequentially in separate experiments: (a) number of hidden layers, (b) dimensions of hidden layers, (c) input layer dropout rate, (d) weight decay, (e) learning rate, and (f) patience. Each boxplot illustrates the performance distribution of the model for the corresponding hyperparameter setting.

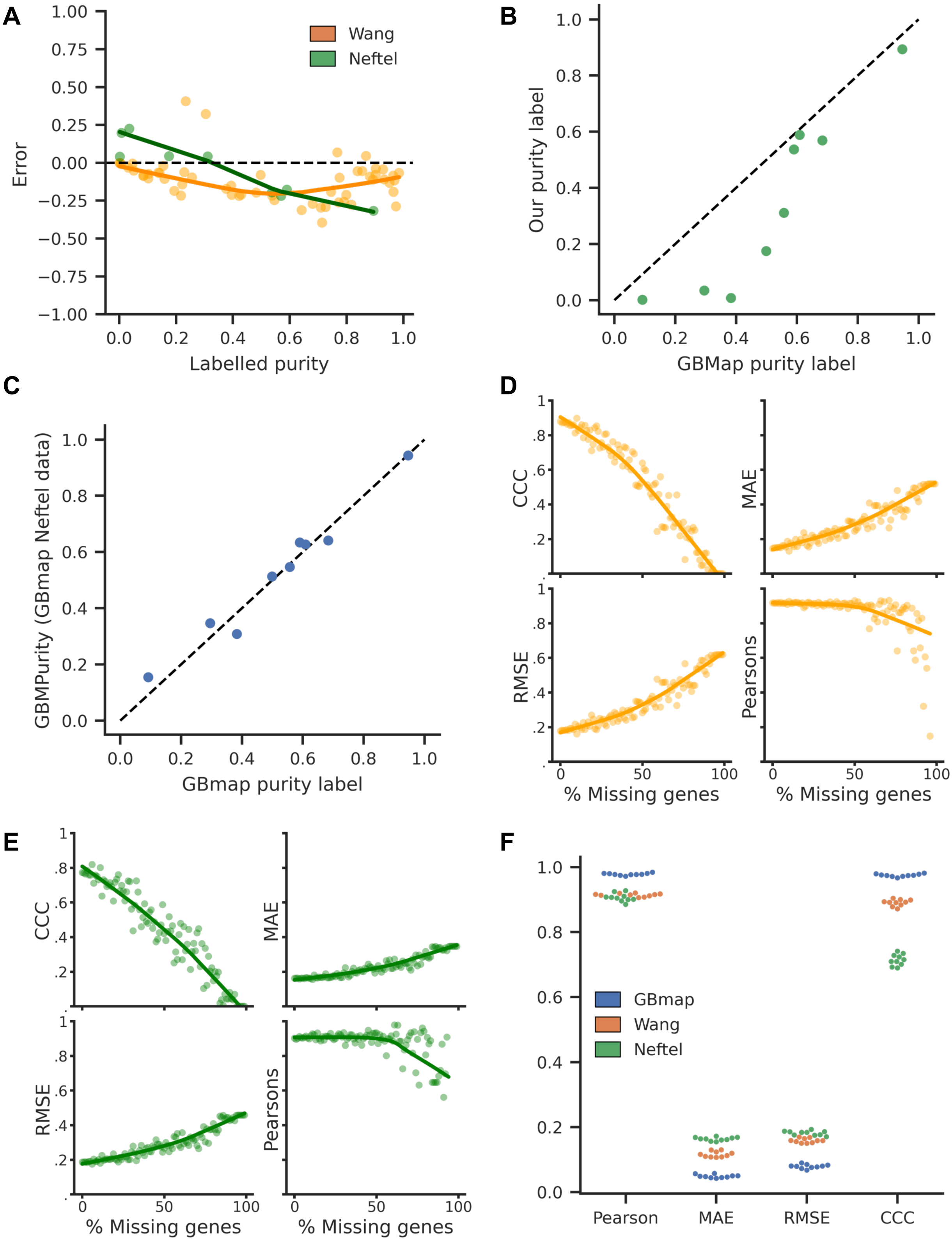


**Supplementary Figure 4. Robustness assessment of GBMPurity.** (a) Scatter plot of GBMPurity prediction error against labelled purity for the Wang et al. (2022) and Neftel et al. (2019) validation datasets. (b) Comparative analysis of Neftel sample purity labels obtained from manual cell labelling versus cell labels from the GBmap resource. (c) Scatter plot depicting GBMPurity predictions on the GBmap Neftel data versus the GBmap purity labels. (d, e) Performance metrics of GBMPurity, including concordance correlation coefficient (CCC), mean absolute error (MAE), root mean squared error (RMSE), and Pearson correlation, evaluated on the Wang and Neftel validation datasets respectively, with varying percentages of randomly zeroed genes. (f) Evaluation of the consistency of 10 different GBMPurity models, each trained with distinct random weight initializations, assessed on the GBmap, Wang, and Neftel pseudobulked samples.

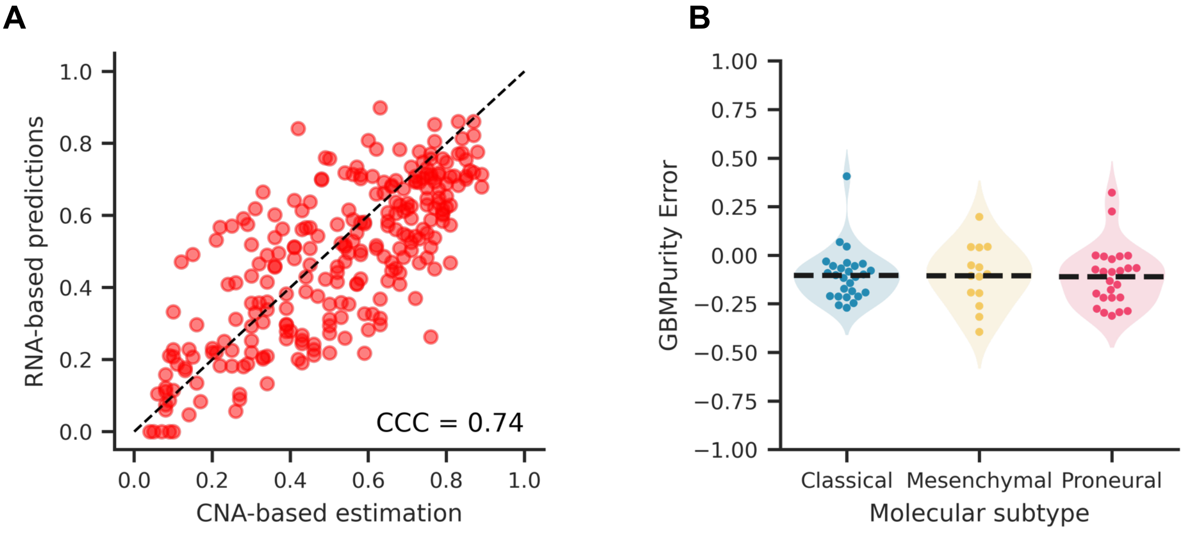


**Supplementary Figure 5. Disagreement between CNA- and RNA-derived purity estimates.** a) GBMPurity was employed to determine the purity of bulk GBM samples obtained from Hoogstrate *et al.*, (2023) (n = 258). These tumour samples underwent both RNA-sequencing and genomic analysis, followed by copy number alteration (CNA) based purity estimation, allowing a direct comparison between our RNA-based method and CNA-based estimates. The plot illustrates the correlation concordance coefficient (CCC) between the two methods. b) Error of GBMPurity p estimates of validation pseudobulks stratified by molecular subtype (n = 66), dashed lines represent population means. ANOVA statistical test revealed no significant difference between groups.
