## Supplementary Table 1 for "GBMPurity: A Machine Learning Tool for Estimating Glioblastoma Tumour Purity from Bulk RNA-seq Data"

Supplementary Table 1. Pairwise comparisons of GBMDeconvoluteR cell scores between bulk GBM molecular subtypes.

| Cell type | A | B | mean(A) | mean(B) | diff | se | p-tukey | p-adjusted |
|---|---|---|---|---|---|---|---|---|
| Astrocyte | Classical | Mesenchymal | 0.355108386 | -0.56692622 | 0.922034607 | 0.089069616 | 1.90E-13 | 8.56E-12 |
| | Classical | Proneural | 0.355108386 | 0.110544073 | 0.244564313 | 0.099283741 | 0.037374773 | 1 |
| | Mesenchymal | Proneural | -0.56692622 | 0.110544073 | -0.67747029 | 0.10543084 | 8.41E-10 | 3.79E-08 |
| Oligodendrocyte | Classical | Mesenchymal | -0.22993251 | -0.13486923 | -0.09506327 | 0.091177747 | 0.550282615 | 1 |
| | Classical | Proneural | -0.22993251 | 0.640496602 | -0.87042911 | 0.101633623 | 1.91E-13 | 8.58E-12 |
| | Mesenchymal | Proneural | -0.13486923 | 0.640496602 | -0.77536584 | 0.107926213 | 6.68E-12 | 3.00E-10 |
| Neuron | Classical | Mesenchymal | 0.0618484 | -0.56648723 | 0.628335629 | 0.086419339 | 3.81E-12 | 1.71E-10 |
| | Classical | Proneural | 0.0618484 | 0.682801289 | -0.62095289 | 0.096329541 | 7.43E-10 | 3.34E-08 |
| | Mesenchymal | Proneural | -0.56648723 | 0.682801289 | -1.24928852 | 0.102293733 | 1.90E-13 | 8.56E-12 |
| Radial glial | Classical | Mesenchymal | 0.0764902 | -0.4465702 | 0.523060403 | 0.091304301 | 4.95E-08 | 2.23E-06 |
| | Classical | Proneural | 0.0764902 | 0.484083853 | -0.40759365 | 0.101774689 | 0.000207637 | 0.009343666 |
| | Mesenchymal | Proneural | -0.4465702 | 0.484083853 | -0.93065406 | 0.108076014 | 1.91E-13 | 8.58E-12 |
| Monocyte | Classical | Mesenchymal | -0.41974891 | 0.938794788 | -1.3585437 | 0.073868896 | 1.90E-13 | 8.56E-12 |
| | Classical | Proneural | -0.41974891 | -0.51180403 | 0.092055122 | 0.082339866 | 0.503348669 | 1 |
| | Mesenchymal | Proneural | 0.938794788 | -0.51180403 | 1.450598822 | 0.087437894 | 1.90E-13 | 8.56E-12 |
| Macrophage | Classical | Mesenchymal | -0.4040508 | 0.796607093 | -1.20065789 | 0.0811601 | 1.90E-13 | 8.56E-12 |
| | Classical | Proneural | -0.4040508 | -0.34076199 | -0.06328881 | 0.090467195 | 0.763824225 | 1 |
| | Mesenchymal | Proneural | 0.796607093 | -0.34076199 | 1.137369083 | 0.096068422 | 1.90E-13 | 8.56E-12 |
| Microglia | Classical | Mesenchymal | -0.36152134 | 0.725284472 | -1.08680581 | 0.084172183 | 1.90E-13 | 8.56E-12 |
| | Classical | Proneural | -0.36152134 | -0.32266419 | -0.03885715 | 0.09382469 | 0.90981671 | 1 |
| | Mesenchymal | Proneural | 0.725284472 | -0.32266419 | 1.047948664 | 0.099633795 | 1.90E-13 | 8.56E-12 |
| DC | Classical | Mesenchymal | -0.40358673 | 0.840416962 | -1.24400369 | 0.079158321 | 1.90E-13 | 8.56E-12 |
| | Classical | Proneural | -0.40358673 | -0.40381743 | 0.000230703 | 0.088235861 | 0.999996231 | 1 |
| | Mesenchymal | Proneural | 0.840416962 | -0.40381743 | 1.244234393 | 0.093698936 | 1.90E-13 | 8.56E-12 |
| Mast Cells | Classical | Mesenchymal | -0.37877187 | 0.845873569 | -1.22464544 | 0.078833081 | 1.90E-13 | 8.56E-12 |
| | Classical | Proneural | -0.37877187 | -0.46003374 | 0.081261873 | 0.087873323 | 0.624846867 | 1 |
| | Mesenchymal | Proneural | 0.845873569 | -0.46003374 | 1.30590731 | 0.093313952 | 1.90E-13 | 8.56E-12 |
| T Cells | Classical | Mesenchymal | -0.42360723 | 0.908349812 | -1.33195704 | 0.075649904 | 1.90E-13 | 8.56E-12 |
| | Classical | Proneural | -0.42360723 | -0.46107748 | 0.037470248 | 0.084325113 | 0.896915283 | 1 |
| | Mesenchymal | Proneural | 0.908349812 | -0.46107748 | 1.369427287 | 0.089546057 | 1.90E-13 | 8.56E-12 |
| NK Cells | Classical | Mesenchymal | -0.41625392 | 0.845984492 | -1.26223841 | 0.078879929 | 1.90E-13 | 8.56E-12 |
| | Classical | Proneural | -0.41625392 | -0.38697034 | -0.02928358 | 0.087925543 | 0.940697583 | 1 |
| | Mesenchymal | Proneural | 0.845984492 | -0.38697034 | 1.23295483 | 0.093369406 | 1.90E-13 | 8.56E-12 |
| B Cells | Classical | Mesenchymal | -0.39987888 | 0.782640693 | -1.18251957 | 0.081767879 | 1.90E-13 | 8.56E-12 |
| | Classical | Proneural | -0.39987888 | -0.32909899 | -0.07077989 | 0.091144671 | 0.717584939 | 1 |
| | Mesenchymal | Proneural | 0.782640693 | -0.32909899 | 1.11173968 | 0.096787844 | 1.90E-13 | 8.56E-12 |
| Plasma B | Classical | Mesenchymal | -0.37609647 | 0.287923285 | -0.66401975 | 0.091639973 | 4.47E-12 | 2.01E-10 |
| | Classical | Proneural | -0.37609647 | 0.326250762 | -0.70234723 | 0.102148855 | 4.97E-11 | 2.23E-09 |
| | Mesenchymal | Proneural | 0.287923285 | 0.326250762 | -0.03832748 | 0.108473346 | 0.933509129 | 1 |
| Mural cell | Classical | Mesenchymal | -0.12689915 | 0.688786542 | -0.8156857 | 0.08247263 | 1.90E-13 | 8.56E-12 |
| | Classical | Proneural | -0.12689915 | -0.72921977 | 0.602320617 | 0.09193024 | 3.86E-10 | 1.74E-08 |
| | Mesenchymal | Proneural | 0.688786542 | -0.72921977 | 1.418006313 | 0.097622052 | 1.90E-13 | 8.56E-12 |
| Endothelial | Classical | Mesenchymal | 0.136699481 | 0.219694943 | -0.08299546 | 0.09235307 | 0.641364348 | 1 |
| | Classical | Proneural | 0.136699481 | -0.57870111 | 0.715400596 | 0.102943726 | 3.08E-11 | 1.39E-09 |
| | Mesenchymal | Proneural | 0.219694943 | -0.57870111 | 0.798396058 | 0.109317432 | 3.09E-12 | 1.39E-10 |
